# Human UFSP1 is an active protease that regulates UFM1 maturation and UFMylation

**DOI:** 10.1101/2022.02.28.482207

**Authors:** David Millrine, Thomas Cummings, Stephen P. Matthews, Joshua J. Peter, Helge Magnussen, Thomas Macartney, Frederic Lamoliatte, Axel Knebel, Yogesh Kulathu

## Abstract

An essential first step in the posttranslational modification of proteins with UFM1, UFMylation, is the proteolytic cleavage of pro-UFM1 to expose a C-terminal glycine. Of the two UFM1-specific proteases (UFSPs) identified in humans, only UFSP2 is reported to be active since the annotated sequence of UFSP1 lacks critical catalytic residues. Nonetheless, efficient UFM1 maturation occurs in cells lacking UFSP2 suggesting the presence of another active protease. We hereby identify a long isoform of UFSP1 to be this protease. Cells lacking both UFSPs show complete loss of UFMylation resulting from an absence of mature UFM1. While UFSP2, but not UFSP1, removes UFM1 from the ribosomal subunit RPL26, UFSP1 acts earlier in the pathway to mature UFM1 and cleave a potential auto-inhibitory modification on UFC1, thereby controlling activation of UFMylation. In summary, our studies reveal important distinctions in substrate specificity and localization-dependent functions for the two proteases in regulating UFMylation.

## Introduction

The ubiquitin-like protein Ubiquitin Fold Modifier-1 (UFM1) is emerging as a central regulator of protein homeostasis through its role in ribosome quality control (Gerakis *et al.*, 2019; Banerjee, Kumar and Wiener, 2020; Witting and Mulder, 2021). The physiological importance of protein UFMylation is evidenced by mutations in UFM1 pathway components that result in neurodevelopmental pathophysiology including cerebellar ataxia, encephalopathy, epilepsy, peripheral neuropathy, and systemic skeletal abnormalities characterised by abnormal cartilage development (Colin *et al.*, 2016; Duan *et al.*, 2016; Egunsola *et al.*, 2017; Di Rocco *et al.*, 2018). In mice, knockout of UFM1 pathway components results in early-stage embryonic lethality that is attributed to defective haematopoiesis (Tatsumi *et al.*, 2011). UFMylation is therefore essential for normal physiology with evidence of involvements in tissue development. Recent studies linking UFM1 to endoplasmic reticulum (ER) stress responses and secretory pathways highlight a potential mechanism to explain these observations. Indeed, UFM1 is conjugated to endoplasmic reticulum (ER)-membrane associated ribosomal subunits (Walczak *et al.*, 2019; Liang *et al.*, 2020) that is induced by ribosomal stalling (Wang et al 2020).

Produced as an 85 amino acid precursor, UFM1 must first be proteolytically activated through the removal of a serine-cysteine dipeptide at its C-terminus (Komatsu *et al.*, 2004). Like ubiquitin, UFM1 is next conjugated to substrates through an enzymatic cascade of E1 (UFM1-Activating Enzyme 5; UBA5), E2 (UFM1-Conjugating Enzyme 1; UFC1), and E3 (UFM1 Specific Ligase 1; UFL1) enzymes, resulting in the formation of an isopeptide bond between the C-terminal glycine of UFM1 and the substrate lysine (Komatsu *et al.*, 2004). These core enzymes are supported by accessory factors that include UFM1 binding protein 1 (UFBP1/DDRGK1) and CDK5 regulatory subunit-associated protein 3 (CDK5RAP3), whose function is poorly defined (Yang *et al.*, 2019; Stephani *et al.*, 2020). UFBP1 and UFL1 localise to the endoplasmic reticulum (ER) and they are thought to catalyse the UFMylation of Ribosomal Protein L26 (RPL26) (Walczak *et al.*, 2019; Stephani *et al.*, 2020). UFMylation of RPL26 in proximity to the Sec61 translocon and oligosaccharyl-transferase (OST) complex occurs after ribosome stalling and initiates specialised autophagy of the ER-membrane to facilitate clearance of potentially toxic misfolded peptides and ribosomes through a lysosomal pathway (Walczak *et al.*, 2019; Liang *et al.*, 2020; Wang *et al.*, 2020). Termed ER-phagy, this organelle-specific degradation pathway involves the wholesale targeting of regions of the rough ER for lysosomal degradation (Khaminets *et al.*, 2015). In a pathway dependent on mitochondrial respiration, acute amino acid starvation stimulates ER-phagy *via* a pathway requiring UFM1 system components and modification of RPL26 (Liang *et al.*, 2020). The UFM1 system may, therefore, control turnover of the translational apparatus in response to cellular and metabolic state. An elaborate system of regulation appears to have evolved solely for this purpose, as RPL26 is the most compelling UFM1 substrate described to date (Walczak *et al.*, 2019).

While the precise function is unclear, it appears that that proper functioning of the pathway requires an equilibrium of UFM1 conjugation that supports stalled ribosome clearance and ER turnover without damaging the cell’s capacity for protein biogenesis. Tight regulation is reflected in the specificity of the pathway which, unlike the highly redundant ubiquitin system, is coordinated by only a handful of enzymes (Komatsu *et al.*, 2004; Gerakis *et al.*, 2019; Walczak *et al.*, 2019; Liang *et al.*, 2020). At present, only two enzymes, UFM1 Specific Protease 1 and 2 (UFSP1 and UFSP2), are known to cleave UFM1 conjugates, with UFSP1 reported to be catalytically inactive in humans (Kang *et al.*, 2007; Ha *et al.*, 2008).

UFM1 requires the peptidase-linked cleavage of the C-terminal Serine^84^-Cysteine^85^ peptide to achieve its mature form, a pre-requisite for the attachment of UFM1 onto substrates. However, in *UFSP2*^−/−^ cell lines, UFM1 modification of the ribosomal subunit RPL26 is enhanced, not abolished (Muona *et al.*, 2016; Walczak *et al.*, 2019), leading us to challenge the notion that UFSP2 is the only UFM1-specifc protease in humans. Hence, we hypothesize that additional unreported mechanisms must exist to support UFM1 maturation in the absence of UFSP2. Indeed, the probable existence of additional UFM1 specific proteases has been noted by others (Walczak *et al.*, 2019; Witting and Mulder, 2021). In the present study we sought to identify this unknown UFM1 peptidase. To our surprise, we isolated UFSP1 from cells that is larger than the presently annotated form and has activity towards the UFM1 precursor. Analysis of knockout cell lines identified overlapping and unique contributions of UFSP1 and UFSP2 to ribosome modification and processing of precursor UFM1. We identify a role for localization of UFSP2 at the ER via its interaction with the ER resident protein ODR4 for its ability to remove UFM1 off RPL26. Further, UFSP1 is unable to remove RPL26 UFMylation highlighting the specificity in the system. Intriguingly, we observe a striking accumulation of UFMylated UFC1 in cells lacking UFSP1. Based on our observations, we propose dual roles for UFSP1 in activating UFMylation, first at the level of UFM1 maturation, and second, by removing a potential autoinhibitory modification on UFC1. Thus, UFSP enzymes act at disparate points of the pathway to ensure appropriate UFMylation.

## Results

### UFSP2 is not the sole UFM1-specific protease in humans

To confirm the existence of additional peptidases with activity towards precursor UFM1, we generated *UFSP2*^−/−^ cell lines using CRISPR-Cas9 approaches. Consistent with previous studies, we observed an increase in both RPL26-(UFM1)1 and RPL26-(UFM1)2 species as a result of UFSP2 deficiency (**Fig-1A**, **Fig-S1A-D**) (Walczak *et al.*, 2019; Liang *et al.*, 2020; Wang *et al.*, 2020; Kulsuptrakul *et al.*, 2021). We next developed an experimental system to monitor UFM1 peptidase activity, where cell lysates were incubated with a reporter protein comprising pro-UFM1^1-85^ fused at its carboxy-terminus to GFP *via* a short peptide linker. Intriguingly, lysates derived from parental wildtype (WT) and *UFSP2*^−/−^ HEK293 cells showed equivalent ability to cleave the UFM1-GFP fusion construct to liberate mature UFM1 suggesting the presence of an additional protease (**Fig-1B**, **Fig-S1E**). Cleavage of the fusion protein could be prevented by pre-incubation with broad-acting cysteine peptidase inhibitors iodoacetmide or N-ethylmaleimide suggesting that the proteolytic activity observed in *UFSP2*^−/−^ cells is due to a cysteine-based protease (**Fig-1C**, **Fig-S1E**). Taken together, these data reveal the existence of additional UFM1-targeting cysteine peptidases with the capacity to activate precursor UFM1.

**Figure-1.**
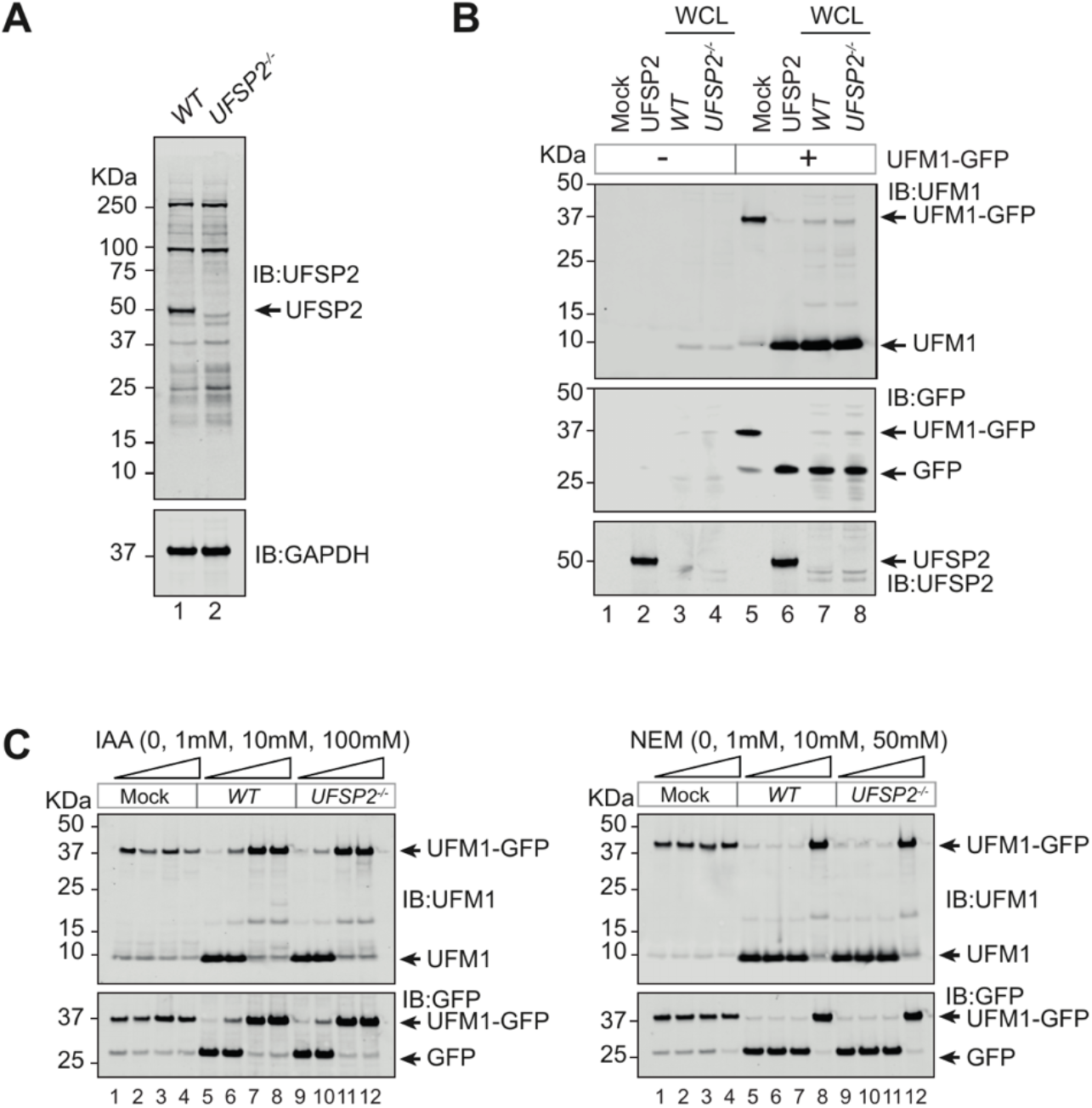
UFSP2 is not the sole UFM1 specific peptidase in human cells. **(A)** Confirmation of UFSP2 knockout by western blotting. Sequencing and details of specific mutations are included in the supplementary materials. **(B)** *In vitro* assay incubating cell lysates from wild type and *UFSP2*^−/−^ HEK293 cells with a UFM1-GFP fusion protein. Cleavage of a recombinant UFM1-GFP fusion protein into its constituent parts, UFM1 and GFP, is interpreted as peptidase activity. Recombinant UFSP2 (2μM) is included as a positive control **(C-D)** Prevention of UFM1-GFP cleavage by the cysteine peptidase inhibitors iodoacetamide (IAA) and *N-ethylmaleimide (NEM).* Cell lysates were pre-treated for 1 hour in the dark at room temperature prior to mixing with the recombinant UFM1-GFP probe. Probe-lysate incubations were performed at 37°C for 2 hours. Data is representative of more than three independent experiments (A-D).

### Isolation of a novel UFM1 peptidase activity from human cells

We next employed classical biochemistry approaches to identify the enzyme responsible. Lysates from HEK293 cells were fractionated sequentially over Heparin and Source-Q columns and ensuing fractions were screened for peptidase activity by monitoring cleavage of the UFM1-GFP reporter (**Fig-2A**). UFM1 specific peptidase activity was restricted to two sequential fractions and strikingly these fractions did not have detectable amounts of UFSP2 (**Fig-2B**, **Fig-S2A**). Importantly, this activity could be recapitulated in fractionations using *UFSP2*^−/−^ cells (**Fig-2C**). We were therefore successful in enriching a novel UFM1 peptidase activity, distinct from UFSP2.

**Figure-2.**
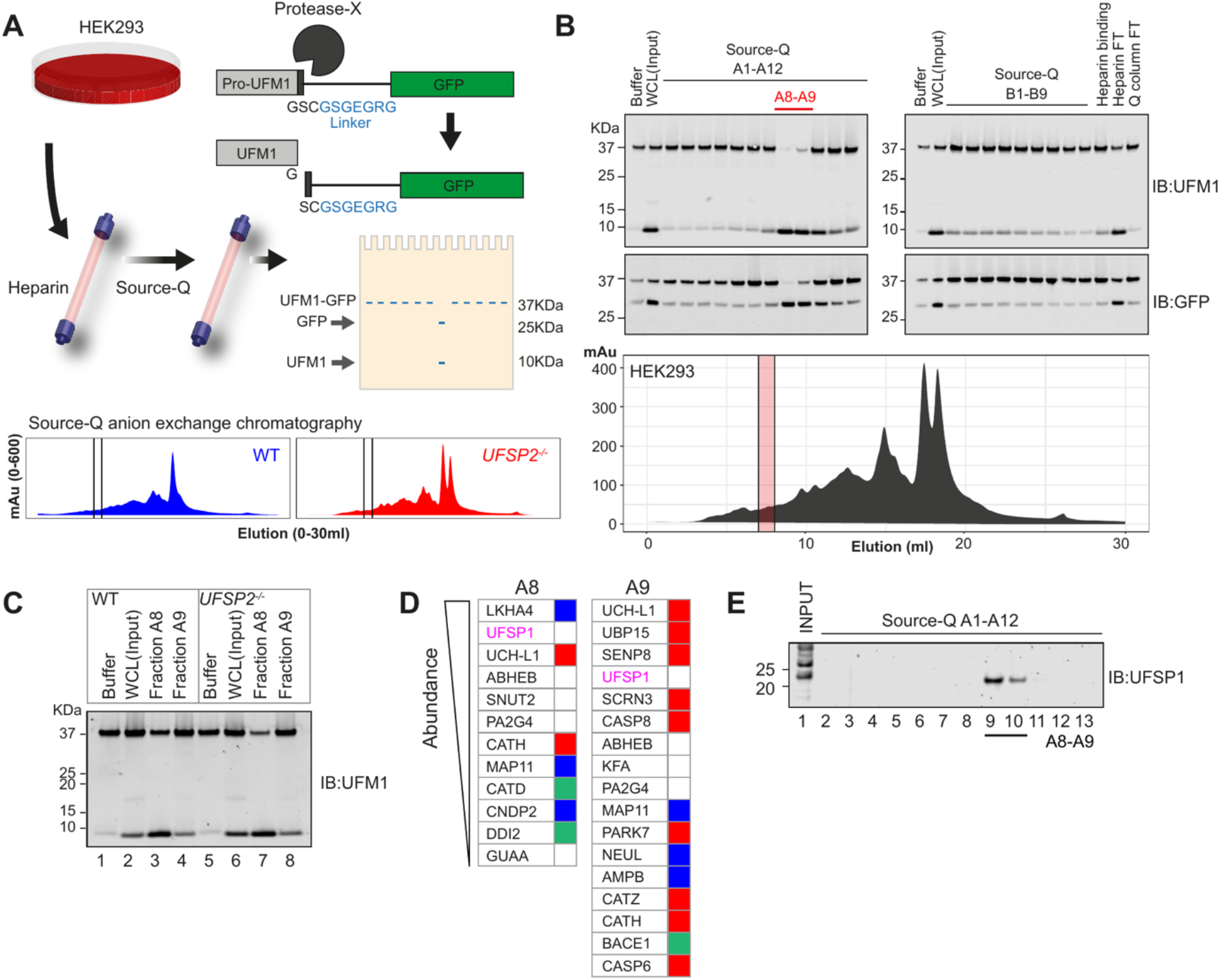
Screening for alternative UFM1 protease identifies UFSP1 as a candidate peptidase. **(A)** (Top) Schematic overview of the screening process. HEK293 cells are lysed by mechanical stress and passed sequentially over Heparin and Source-Q columns by FPLC. Eluted fractions are screen for activity by incubation with the UFM1-GFP fusion protein. (Below) Chromatograms showing protein eluted from Source-Q columns on a salt gradient. Shown are experiments performed in parallel using unmodified HEK293 wild type (blue) and *UFSP2*^−/−^ (red) HEK293 cells **(B)** Representative screening results. Heparin binding protein has been eluted in a single fraction (‘Heparin binding’) while Source-Q binding protein is eluted in fractions A1-B9. Eluted fractions were incubated with UFM1-GFP fusion protein for 2 hours at 37°C and analysed by immunoblot. Heparin and Q-column Flow-Through (FT) are shown far right. Activity is detected in Q column fractions A8 and A9 (red lines). **(C)** *In vitro* assay incubating active fractions (A8 and A9) purified from unmodified and *UFSP2*^−/−^ HEK293 cell lysates with the UFM1-GFP fusion protein. Experiment is the same as depicted by chromatograms in (A). (D) Identity of proteases identified in the active fractions. The output of Mass-spectrometry analysis of the active fractions was aligned with MEROPS database annotations (data frames were merged in R studio by gene symbol) to identify proteases. **(E)** Immunoblot analysis of endogenous UFSP1 in active fractions.

To identify the protease, we adopted an unbiased approach using LC-MS profiling of the active fraction from *UFSP2*^−/−^ cells. These data identified 974 proteins including 21 proteins documented as having deubiquitinase or hydrolase activity (**Fig-S2B**). Among these, Ubiquitin Carboxy-terminal Hydrolase L1 (UCHL1) distributed in close alignment with the novel peptidase activity (**Fig-S2C**), an attractive candidate given its historic association with the maturation of ubiquitin precursors (Grou *et al.*, 2015). However, when tested, neither recombinant UCHL1 nor related family members (UCHL3, UCHL5 or BAP1) could cleave the UFM1-GFP reporter (**Fig-S2D**). Hence, we depleted UCHL1 using CRISPR-Cas9 and repeated the fractionation and mass spectrometry analyses (**Fig-S2E-G**). To our surprise, these analyses revealed UFM1 specific peptidase-1 (UFSP1), characterised as an inactive homologue of UFSP2, among the top candidates in both active fractions (**Fig-2D**). Analysis by immunoblotting confirmed the restricted distribution of UFSP1 in the two active fractions (**Fig-2E**). Therefore, our cell-fractionation studies successfully captured an active form of UFSP1, a surprising observation considering the present consensus surrounding UFSP1 non-functionality in humans (HGNC annotation).

### Human UFSP1 is an active cysteine protease

The UFSP1 activity we observe raises the possibility that the HGNC annotation is incorrect and the UFSP1 expressed in cells spans a longer stretch at the N-terminus that contains the catalytic residues. Aggregate ribosome profiling data across multiple studies (Ribo-seq; GWIPS-viz; https://gwips.ucc.ie/) supported our hypothesis that regions upstream of the annotated *UFSP1* locus are actively translated. A high number of protected reads was observed 5’ to the incorrect start site with coverage of the catalytic cysteine (C54) and a putative CTG start codon (**Fig-3A**). Reported previously, this non-canonical initiation site is a rare example of translation initiation from codons other than ATG and is proposed to be the only in-frame start codon capable of producing a functional UFSP1 cysteine protease (Ivanov *et al.*, 2011). We therefore re-analysed mass-spectrometry data from the two active fractions to search for peptides that correspond to regions upstream of the annotated start site. This analysis identified 12 matching peptides corresponding to the N-terminus of UFSP1 (65% coverage) and included the catalytic cysteine (**Fig-3B**). Indeed, close inspection of curated isoforms of UFSP1 (Uniprot, Ensembl, and NCBI) showed that of the two human UFSP1 variants, one isoform (A0A5F9ZGY7) shares amino acid residues described as essential for the catalytic activity of murine UFSP2 (**Fig-3C**) (Kang *et al.*, 2007; Ha *et al.*, 2008). This conclusion is supported by cross-species bioinformatic analysis of *UFSP1* transcripts that support the existence of the longer version (**Fig-S3A-C**). Further confirmation is obtained in immunoblotting of endogenously expressed UFSP1 which migrates at a size consistent with the predicted molecular mass (~24kDa) upon translation of the correct transcript (**Fig-2E**). Of note, this is larger than the incorrectly HGNC annotated UFSP1 (15kDa).

**Figure-3.**
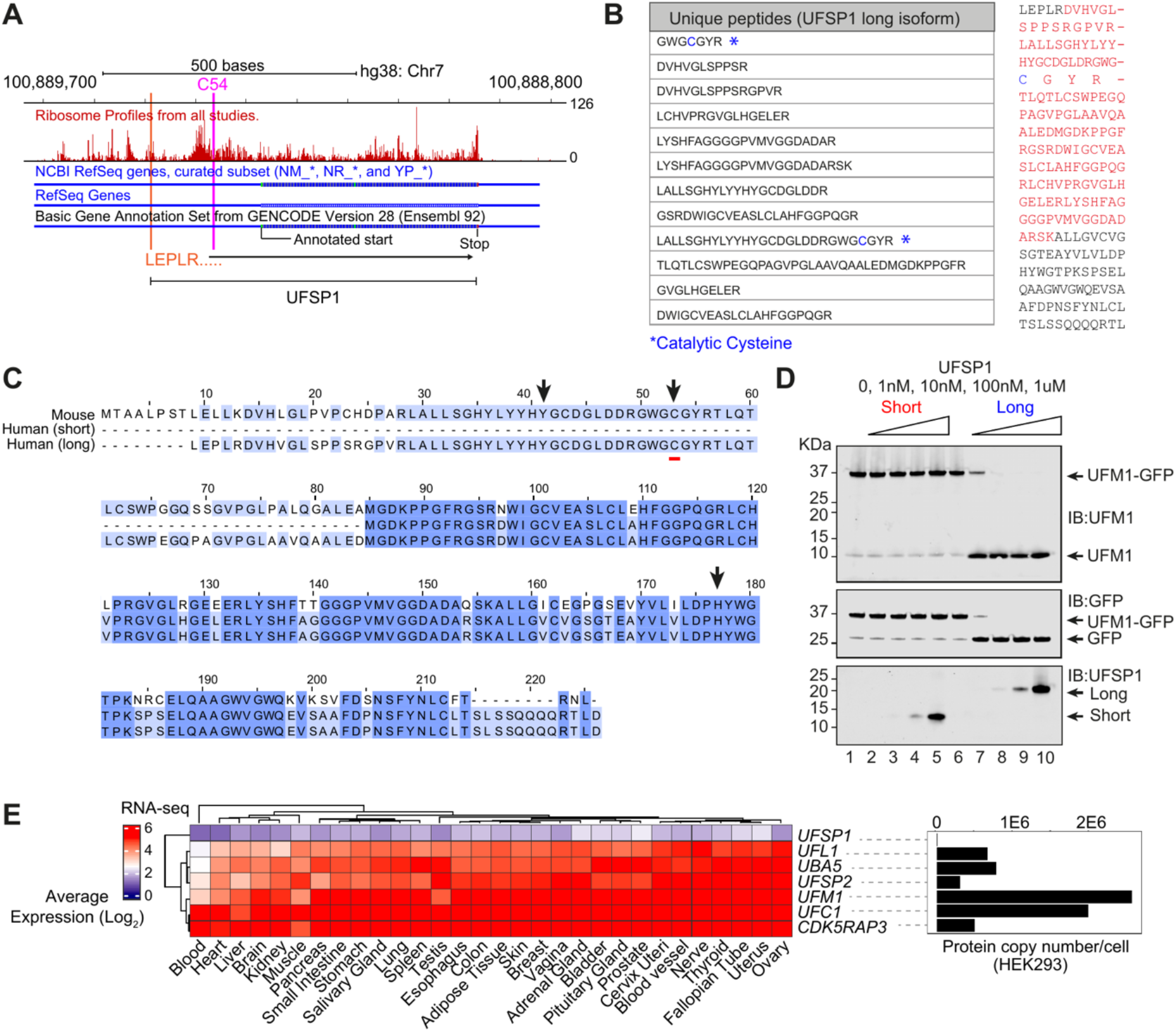
UFSP1 is an active protease. **(A)** Ribosome profiling data downloaded from GWIPS-viz (https://gwips.ucc.ir/). Gobal aggregate protected reads from multiple studies are shown in red (UCSC-track) aligned to Refseq and GENCODE (v28) gene annotation. **(B)** (left) LC/MS data showing peptides mapping to long isoform UFSP1 identified in active fractions. (right) Proposed full-length UFSP1. Sequences in red are identified in LC/MS analysis of active fractions. The catalytic cysteine is highlighted blue. (**C)** Cross-species multiple sequence alignment of UFSP isoforms. Human short (Q6NVU6), human long (A0A5F9ZGY7), and mouse (Q9CZP0) UFSP1. **(D)** *In vitro* assay incubating recombinant long and short isoform UFSP1 variants with the UFM1-GFP fusion protein. Details on expression and purification are included in the supplementary materials. **(E)** (left) RNA-sequencing analysis of human tissues by the Genotype Tissue (GTEx) consortium (https://gtexportal.org/home/). Shown is the Log2 transformed Transcripts Per Million (TPM). (right) Data Independent Acquisition (DIA) quantitative proteomics analysis of UFM1 pathway components in the HEK293 cell line.

These results imply that the long isoform is the primary UFSP1 protein expressed in human cells that is catalytically active. To test this experimentally, we expressed and purified recombinant UFSP1 corresponding to the 24kDa form and the incorrectly annotated protein (**Fig-S3D**). When incubated with the UFM1-GFP fusion protein, only the recombinant long-isoform UFSP1, but not its truncated form, showed cleavage activity *in vitro* (**Fig-3D**). Of note, UFSP1 is expressed at very low levels in cells as judged by RNA-sequencing data (GTEx) and quantitative proteomics data (**Fig-3E**), possibly explaining why the correct UFSP1 may have escaped detection further contributing to the acceptance of the misannotated form. Taken together, our analyses provide convincing evidence that the long isoform UFSP1 we here identify is the correct endogenous UFSP1 that is catalytically competent *via* a cysteine-thiol based mechanism and corrects the long-held misconception that human UFSP1 is an inactive protease.

### Characterization of UFSP1

Since nothing is known about the function of UFSP1 in cells, we next sought to identify the roles of UFSP1 in regulating UFMylation and to dissect the relative contributions of UFSP1 and UFSP2. First, a comparison of UFMylation in *UFSP1*^−/−^ and *UFSP2*^−/−^ cells revealed an increase of RPL26 UFMylation, which was missing in the WT and *UFSP1*^−/−^ cells. Instead, *UFSP1*^−/−^ cells showed an increase in UFMylated UFC1 that could be confirmed in immunoprecipitations of UFM1 (**Fig-4A**). To confirm these observations and identify the lysine residue on UFC1 that is modified we immunoprecipitated UFM1 from *UFSP1*^−/−^ cell lines and analysed by LC-MS. Importantly the immunoblotting data were corroborated by detection of MS spectra corresponding to UFC1 peptides in which Lys122 was modified by Val-Gly (the C-terminal UFM1 dipeptide that remains on UFMylated residues after tryptic digest) (**Fig-S4A**). Hence, in the absence of UFSP1, there is an accumulation of UFC1 UFMylated at K122. Interestingly, K122 is situated near the catalytic cysteine of UFC1 (**Fig-4B**).

**Figure-4.**
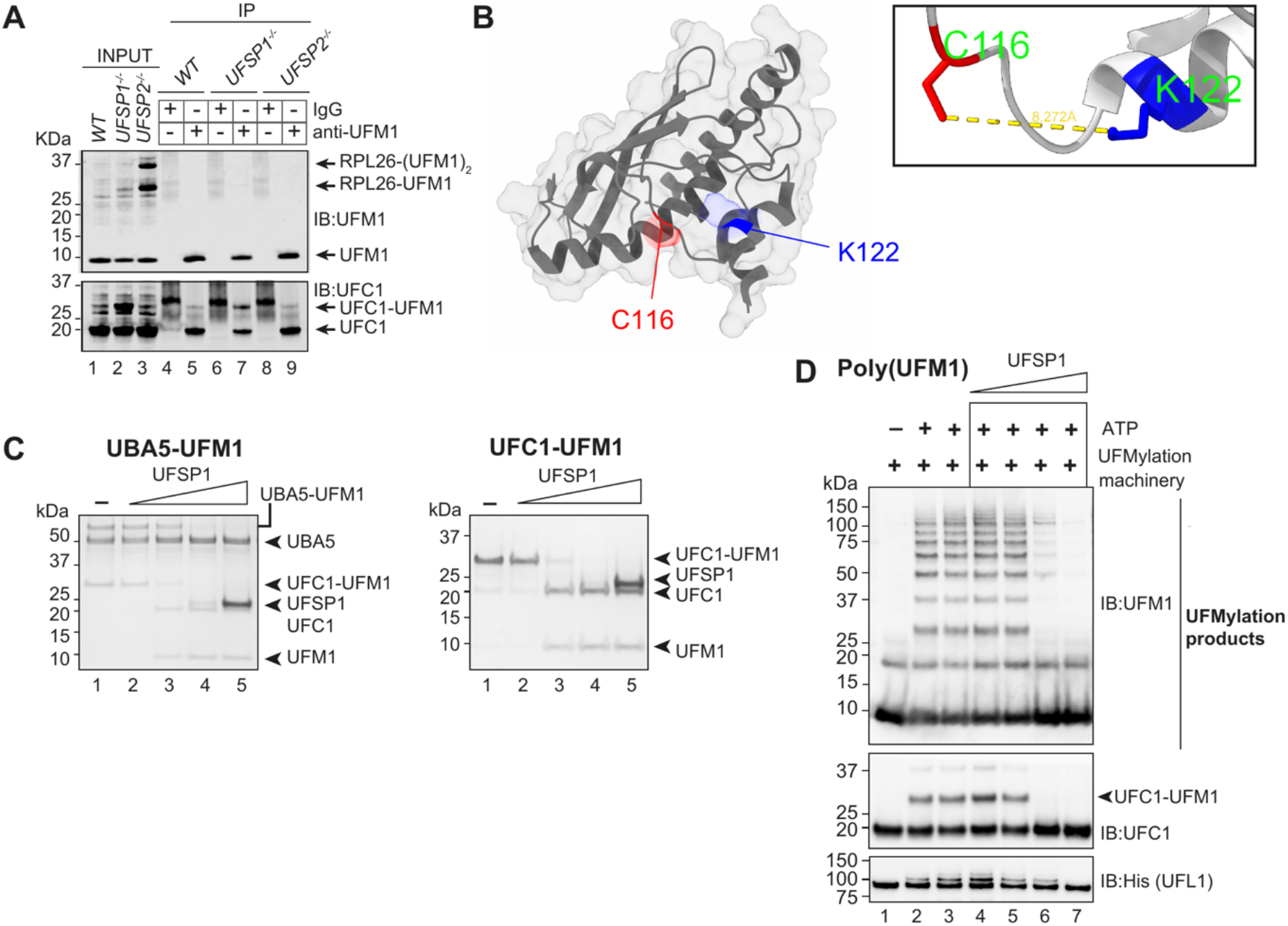
UFSP1 is active against diverse substrates in vitro. **(A)** Immunoprecipitation of UFM1 from the indicated cell lysates **(B)** Crystal structure of UFC1 (pdb2z60) with K122 (blue) and C116 (red) highlighted. Distance between residues (8.272A was calculated using ChimeraX. **(C)** Activity of recombinant UFSP1 against the indicated substrates. **(D)** Activity of UFSP1 against UFMylation products. UFM1 pathway components were reconstituted *in vitro* (UBA5, UFC1, UFBP1-UFL1) in the presence of ATP. After 1 hour the reaction was quenched with Apyrase and incubated with increasing molar concentrations of recombinant UFSP1. Note the dose-effect disassembly of poly (UFM1) chains.

While the earlier assays used a UFM1-GFP reporter to reveal peptidase activity in UFSP1, to confirm that USFP1 has isopeptidase activity, we performed a series of *in vitro* assays with various *in vitro* generated substrates including UBA5 and UFC1 modified with UFM1 *via* an isopeptide bond. Further, through the reconstitution of UFM1 pathway components, we were able to synthesise UFMylated products (Peter JJ et al, *manuscript in preparation*). These are a heterogenous mixture of UFM1 conjugates including K69-linked polyUFM1 chains in addition to auto-modified UFC1 and UFL1 (Peter JJ et al, *manuscript in preparation*). UFSP1 effectively cleaved UFM1 from these different substrates and catalysed the disassembly of polyUFM1 chains *in vitro* (**Fig-4C-D**). Taken together, these data show that in addition to its peptidase function in targeting precursor UFM1, UFSP1 is an effective isopeptidase with activity towards diverse substrates.

### UFSP1 and UFSP2 act at separate points in the UFM1 pathway

To further explore the observation of enriched K122-modified UFC1 and establish that these observations are not cell line specific, we generated a series of *UFSP1* and *UFSP2* knockouts in three different cell lines, HEK293, U2OS, and HeLa, and multiple clones were confirmed by sequencing (**Fig-S5A**). Consistent with our earlier observation (**Fig-S1C-D**), we observe increased mono and di-UFMylated RPL26 in *UFSP2*^−/−^ but not *UFSP1*^−/−^ cell lines (**Fig-5A**). Meanwhile, in all the different *UFSP1*^−/−^ cell lines tested, we observed a size shift in UFC1 of approximately 10KDa (**Fig-5A**). This size shift corresponded to modification of UFC1 with UFM1 and accounted for up to 50% of cellular UFC1 in *UFSP1*^−/−^ cell lines (**Fig-5A**). These results suggest that UFC1 is constitutively modified with UFM1 in cells whose removal depends on UFSP1. In the absence of either UFSP1 or UFSP2, cells appear able to generate sufficient mature UFM1. However, in cell lines lacking both UFSP enzymes (*UFSP1*^−/−^/*UFSP2*^−/−^), a complete loss-of-function phenotype manifested by a total absence of detectable UFMylation (**Fig-5B**). These data are consistent with a requirement for either UFSP1 and/or UFSP2 in the generation of mature UFM1 and suggest that both enzymes contribute, in a partially redundant manner, to precursor UFM1 maturation. To confirm that the complete loss of UFMylation observed in double knockout cell lines stems from an absence of mature UFM1 in these cells, *UFSP1*^−/−^/*UFSP2*^−/−^ cell lines in HEK293 and HeLa backgrounds were transiently transfected with constructs expressing either mature UFM1^(1-83)^ or its precursor counterpart (UFM1^1-85^). Over-expression of mature, but not proUFM1, successfully rescued mono and di-UFMylated RPL26 (**Fig-5B**). These data are consistent with *in vitro* analysis of peptidase activity in cell lysates where cell lysates from *UFSP1*^−/−^, *UFSP2*^−/−^, *UFSP1*^−/−^/*UFSP2*^−/−^ HEK293 cell lines were incubated with the proUFM1-GFP probe. Here, cleavage activity was only completely abolished in the absence of both enzymes (**Fig-5C**). Further, close inspection of immunoblot analyses reveals a size shift in UFM1 migration consistent with an increase in molecular weight corresponding to proUFM1 in cell lines lacking both UFSP enzymes (**Fig-5D**). While these experiments clearly demonstrate complete loss of UFM1 maturation in cells lacking both UFSP1 and UFSP2, this does not preclude the existence of additional proteases in specific cell types.

**Figure-5.**
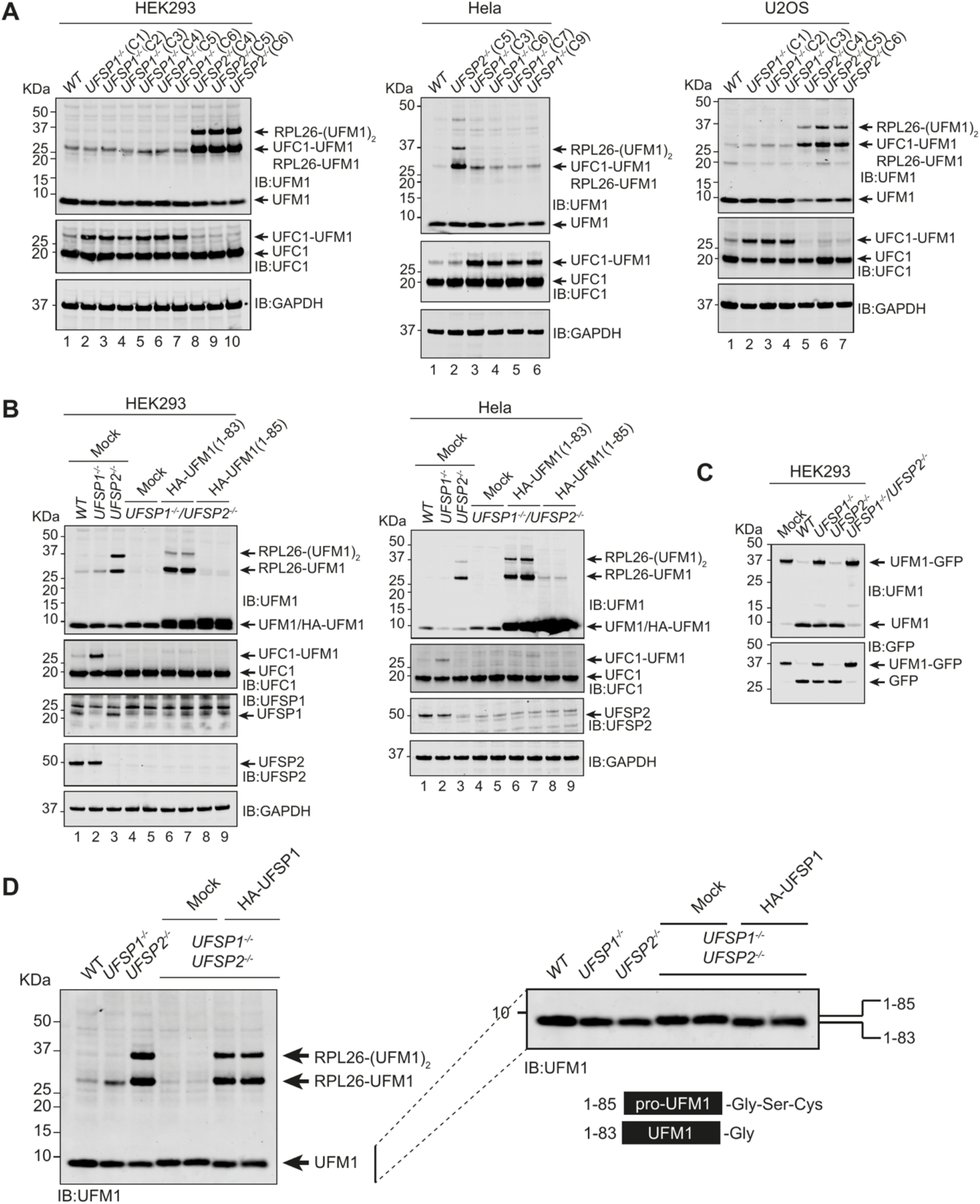
UFSP1 is the UFM1 activating peptidase *in vivo*. **(A)** Immunoblot analysis of *UFSP1*^−/−^ and *UFSP2*^−/−^ cell lines as indicated **(B)** Rescue of RPL26 UFMylation by expression of mature UFM1. Constructs expressing HA-tagged mature (UFM1^1-83^) or precursor (UFM1^1-85^) UFM1 were transiently transfected into UFSP1^−/−^/UFSP2^−/−^ double knockout cell lines. 24 hours later cells were lysed and analysed by immunoblot. **(C)** *In vitro* assay incubating HEK293 cell lysates from the indicated knockout cell lines with the UFM1-GFP probe. **(D)** Immunoblot analysis of HEK293 cells transiently transfected with HA-tagged UFSP1. (right) zoomed in section of the blot shown left to highlight changes in UFM1 migration. This western blot is reproduced complete with loading controls in the supplementary materials (**Fig-S5B**). Labels in (A) include an abbreviated Clone ID (e.g., C1 is Clone-1) for tracking to the MRC-PPU cell bank. Cell lines have been sequenced with representative sequencing traces included in the supplementary materials.

### Subcellular localization regulates function of UFSPs

We next explored whether differences in subcellular localization of UFSP1 and UFSP2 might contribute to the differences in substrate specificities observed in knockout cells. UFSP2 interacts with ODR4 (Odorant Response Abnormal protein-4), a transmembrane protein that is localised to the ER membrane where it is thought to anchor UFSP2 in proximity to the ER-ribosome interface (Chen *et al.*, 2014). In contrast, sequence analysis and structural predictions suggest that UFSP1 will not interact with ODR4 and is likely to instead reside in the cytosol. This makes it more possible for UFSP1 to be the protease mainly responsible for UFM1 maturation. To disrupt the ER localization of UFSP2, we generated ODR4 knockout cell lines and monitored UFMylation (**Fig-6A**, **Fig-S6A**). Intriguingly, UFSP2 protein levels are markedly reduced in *ODR4*^−/−^ cell lines, and vice-versa, suggesting that UFSP2 and ODR4 are engaged in a mutually stabilising relationship (**Fig-6A**). Moreover, UFSP2 and ODR4 knockout cell lines were an exact phenocopy in their role in restraining levels of RPL26 UFMylation (**Fig-6A**). In cell lines lacking both UFSP1 and ODR4, levels of di-UFMylated but not mono-UFMylated RPL26 were reduced. Interestingly, this is similar to our observation of a cell line heterozygously deficient for UFSP1 (*UFSP1^+/-^*/*UFSP2^−/−^)* where the second UFM1 modification on RPL26 was absent (**Fig-6A**, **Fig-S6B**). These data may indicate a preference for mono-UFMylated RPL26 in circumstances where mature UFM1 is limiting and aligns with the description of RPL26 modification as sequential with K132 UFMylation dependent on prior modification of K134 (Walczak *et al.*, 2019). Finally, we investigated whether UFSP1 could contribute directly to the UFMylation pathway in ribosomal quality control. Upon induction of ribosome stalling following treatment with low dose anisomycin, RPL26 UFMylation is induced in the membrane fraction of WT cells (**Fig-6B**, top panel). In the absence of UFSP2, significant RPL26 UFMylation is observed which does not increase further upon ribosome stalling (**Fig-6B**, bottom panel). In contrast, UFSP1 knockout cells remained competent for the induction of ribosome UFMylation upon ribosome stalling (**Fig-6B**). These data suggest that ribosome UFMylation is dynamic and is mainly regulated by UFSP2. UFSP1 on the other hand indirectly regulates RPL26 UFMylation by facilitating UFM1 maturation and cleaving UFM1 from UFMylated UFC1.

**Figure-6.**
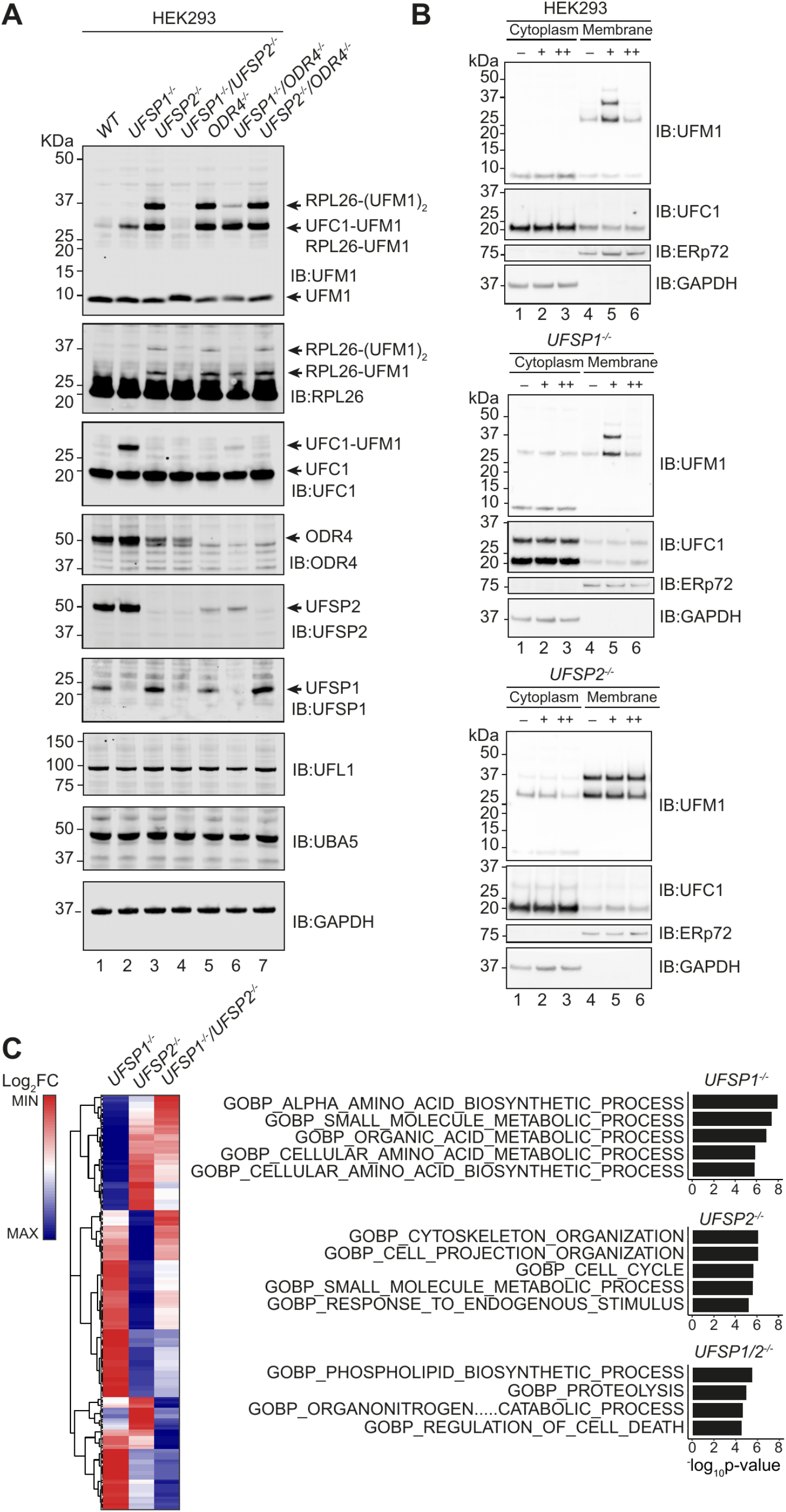
Schematic overview of the UFM1 system. **(A)** Immunoblot analysis of the indicated knockout cell lines (HEK293 Flp-in TREx). Sequencing data is shown, in part, in the supplementary materials. Full data is available on request from the study lead author. **(B)** Inducible UFMylation. Cells lines were treated with 200nM (+) or 50μM (++) Anisomycin for 20 minutes prior to cell lysis. Immunoblot analysis of cytoplasmic and membrane fractions is shown. Note the cytosolic localisation of UFC1, while UFMylated RPL26 localises to the membrane fraction. **(C)** (left) heatmap showing proteins passing statistical thresholds in at least one experimental condition (Benjamini & Hochberg adjusted p-value <0.05; Log2FC>1). Colouring is the scaled Log2 Fold Change by row. (right) Gene Ontology (Biological Processes) enrichments calculated using a hypergeometric distribution test (Broad Institute; http://www.gsea-msigdb.org/gsea/msigdb/compute_overlaps.jsp). Heatmap is hierarchically clustered by the Euclidean method.

To address the biological significance of these findings, the proteomes of the different cell lines (*UFSP1*^−/−^, *UFSP2*^−/−^, *UFSP1*^−/−^/*UFSP2*^−/−^ HEK293) were analysed by Data Independent Acquisition (DIA) quantitative proteomics. A total of 191 proteins passed the fold change and significance thresholds in at least one phenotype (Benjamini & Hochberg adjusted p-value <0.05; Log2FC>1) **(Fig-6C, Fig-S6C)**. Heatmap analysis (hierarchical clustering by the Euclidean method) revealed that distinct subsets of proteins are altered in UFSP1 and UFSP2 knockout cells (**Fig-6C**, **Fig-S6D-F**). Gene Ontology overlaps, calculated using a hypergeometric distribution tool provided by the molecular signatures database (msigdb; https://www.gsea-msigdb.org/), suggested contributions of UFSP1 to small molecule metabolism (**Fig-6C, Fig-S6F**). By contrast, in *UFSP2*^−/−^ HEK293 cells, processes including metabolism (IRS2), cytoskeletal remodelling (RHOB), and cellular motility/morphology (CDK10, APOE, WASL, PRKG1) were heavily represented. *UFSP1*^−/−^/*UFSP2*^−/−^ cells exhibited an intermediate phenotype with features of both *UFSP1*^−/−^ and *UFSP2*^−/−^ knockout cell lines (**Fig-6C**, **Fig-S6C-F**).

## Discussion

Considering that 99 DUBs have been identified in the analogous ubiquitin system (Lange, Armstrong and Kulathu, 2021), it seemed remarkable that the UFM1 pathway in humans could be reliant on only one enzyme. Together with the confounding observation of active UFMylation in *UFSP2*^−/−^ cell lines, it has been clear to many in the field that additional enzymes must exist to process precursor UFM1 into its mature counterpart (Witting and Mulder, 2021). Our study now reveals that the annotation of UFSP1 as inactive is mistaken. We report that human UFSP1 is not only an active cysteine protease that is expressed in cells, but also one with key functions in the UFM1 pathway. Overall, we define at least three contributions of UFSP enzymes. First, consistent with reports elsewhere, we find UFSP2 to be a key regulator of RPL26 modification. Second, UFSP1 acts to restrain levels of UFC1 modified with UFM1. Finally, in a partially redundant manner, both UFSP enzymes contribute to UFM1 precursor maturation and the maintenance of a cellular pool of mature UFM1 (**Fig-7A**).

One of two mechanisms may contribute to the different substrate activities of UFSP enzymes. First, the cellular expression of levels of UFSP1 are very low, possibly limiting its ability to counter RPL26 UFMylation. Second, cellular localisation. The cytosolic localization of UFSP1 compared to the ER localization of UFSP2 makes it more likely for UFSP1 to be the UFM1 maturing enzyme. The transmembrane domain of ODR4 provides a means for UFSP2 to associate with the ER-membrane, bringing it into proximity of the ribosome and UFM1 ligase machinery (Chen *et al.*, 2014). Structure prediction of ODR4 using AlphaFold reveals it to adopt an MPN fold (Jumper *et al.*, 2021). Interestingly, the crystal structure of Ufsp from *C.elegans* reveals the presence of an MPN fold in addition to the catalytic domain (Kim, Ha and Kim, 2018). Here, it remains to be investigated whether ODR4 allosterically modulates the activity of UFSP2. Superposition of an Alphafold predicted complex of UFSP2-ODR4 reveals that UFSP1 lacks domains required for interaction with ODR4 (**Fig 7B**). Hence, UFSP1 may be localised elsewhere, likely the cytosol where it is unable to interact with UFMylated RPL26 at the ribosome.

**Figure-7.**
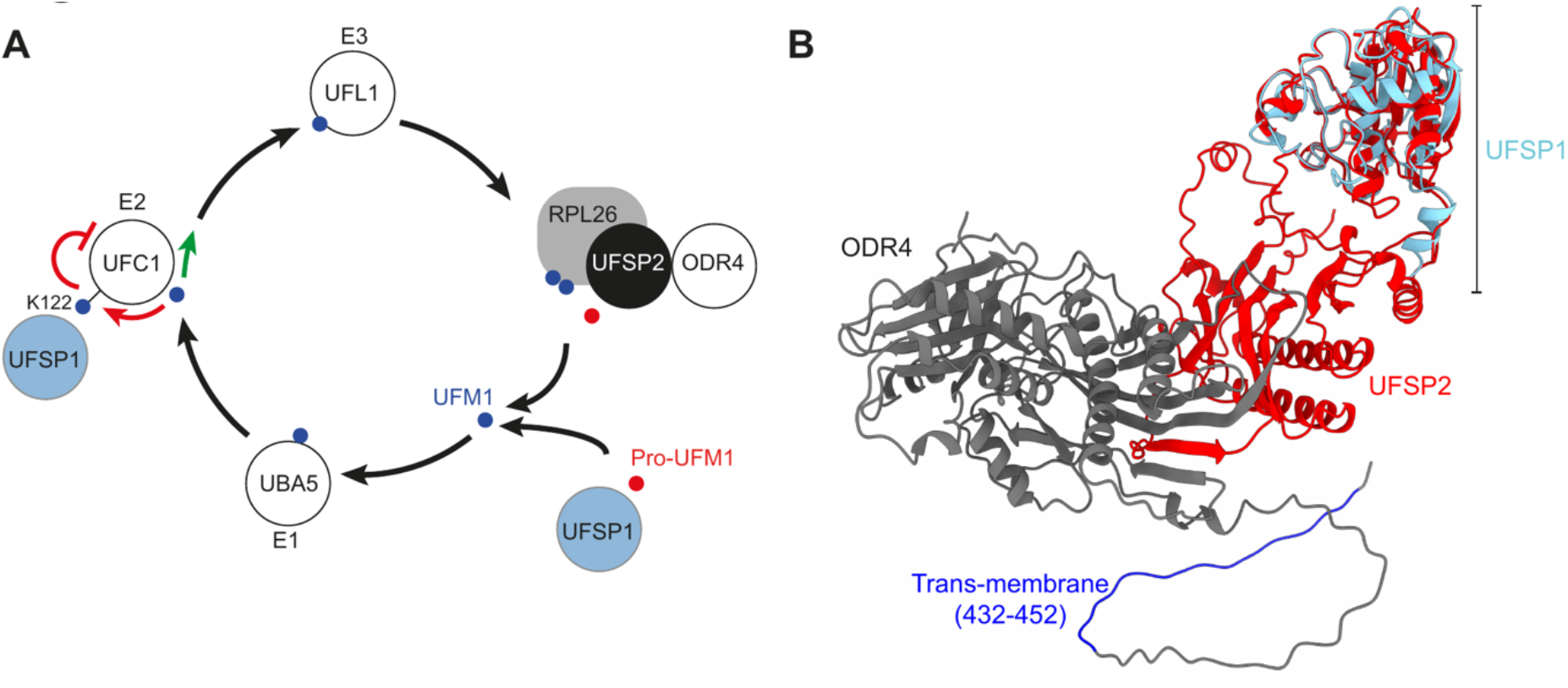
**(A)** Schematic showing suggested model of the UFM1 pathway. Briefly, precursor UFM1 is proteolytically activated through the removal of a C-terminal Serine-Cysteine peptide prior to sequential loading onto the E1, E2, and E3 conjugating enzymes. This culminates in modification of the ribosomal subunit RPL26. UFSPs act at several points in this pathway; (1) Both UFSP1 and UFSP2 contribute to pro-UFM1 processing; (2) UFSP1 catalyses the removal of UFM1 from UFC1, releasing UFC1 from a potentially auto-inhibitory state; (3) UFSP2 catalyses the removal of UFM1 from RPL26, preventing excess ribosome modification. ODR4 is essential for stabilising UFSP2 and anchoring it the ER membrane in proximity to the ribosome. **(B)** Alphafold prediction of UFSP2-ODR4 complex aligned to the predicted structure of human UFSP1.

We observe a striking accumulation of UFMylated UFC1 in *UFSP1*^−/−^ cells but not *UFSP2*^−/−^ cells. This suggests that UFC1 is constantly UFMylated in cells and this modification is removed by UFSP1. Interestingly, previous studies have suggested regulation of E2 activity by covalent modification (Stewart *et al.*, 2016). It may be that UFC1 UFMylation alters protein interactions with other UFM1 conjugating enzymes or putative substrates, either sterically or through the addition of a UFM1-interacting surface. For instance, the SUMO-specific E2 Ubc9 (UBE2I) is auto-SUMOylated, a modification that serves to attract proteins with SUMO interacting motifs (SIMs) (Stewart *et al.*, 2016). Of note, a UFM1 interacting motif has been described and may serve a similar purpose here (Padala *et al.*, 2017; Kumar *et al.*, 2021). Alternatively, auto-UFMylation may interfere with E2 catalytic activity. A recent case study of UBE2S auto-modification identified a lysine residue precisely five amino acids from the catalytic cysteine (K^+5^) (Liess *et al.*, 2019). In thioester transfer assays, auto-modification at this site on UBE2S reduced the overall efficiency of ubiquitin transfer from the E1 by steric hindrance. Intriguingly, up to 25% of E2 enzymes possess a conserved lysine residue at +5 amino acids form the catalytic cysteine (Liess *et al.*, 2019). Following a similar theme, monoubiquitylation of UBE2T at K86 (5 amino acids from the catalytic cysteine K91) is suggested to reduce E2 activity linked to the Fanconi Anemia DNA repair pathway mediated by the E3 ubiquitin ligase FANCL (Machida *et al.*, 2006). Our work identifies UFC1 to be modified on K122, a site located +6 residues from the catalytic C116. While further work is required, we speculate a similar inhibitory role for this UFC1 modification. These observations may reflect a common regulatory feature of E2 enzymes. If so, then UFSP1 may the first documented instance of a protease acting to relieve E2 autoinhibition, eventually influencing the rate of overall UFMylation.

Hence, UFSP1 may act at two levels to activate UFMylation, firstly in UFM1 maturation and secondly as part of a regulatory loop to remove the inhibitory UFC1 modification. Given that *UFSP1* gene expression is remarkably low relative to other UFM1 pathway components, it is possible that under specific circumstances, *UFSP1* gene induction may function as an inducible ON-OFF switch or accelerator for ribosome UFMylation. Further studies will be required to assess whether this is the case and to assess the precise kinetics of UFMylation in cellular models with *UFSP1* over-expression or deletion. Moreover studies examining if UFSP1 contributes to other functions attributed to the UFM1 pathway including regulation of the DNA damage response (DDR) *via* histone H4 UFMylation, or ER-phagy induced by metabolic stress is likely to reveal novel insights (Qin *et al.*, 2019; Liang *et al.*, 2020).

A serendipitous outcome of our study is correction of the long-held misconception of UFSP1 as an inactive peptidase homolog. This misunderstanding stems from the annotation of UFSP1 as the ‘inactive UFM1 specific protease-1’ (HUGO Gene Nomenclature Committee (HGNC)) and is based on a hypothetical interpretation of an N-terminal truncated version of UFSP1 that lacks catalytic residues when compared with UFSP2. Perhaps early studies characterising UFSP1, performed in murine systems, were reliant on annotations that pre-date entry of the long isoform described here (Kang *et al.*, 2007; Ha *et al.*, 2008). Clearly, the naming of UFSP1 as the ‘Inactive UFM1 specific protease-1’, will require revision. Our study corrects the view that human UFSP1 is catalytically inactive, and in doing so, has laid foundations for future investigation into the unique contributions of UFSP family proteases to ER and cellular homeostasis.

## Acknowledgements

The authors would like to thank Ron Kopito and his lab for helpful discussions and input, support staff at MRC-PPU reagents and services. This study was funded by an ERC starting grant (RELYUBL, 677623), MRC grant MC_UU_00018/3, BBSRC (BB/T008172/1) and the Lister Institute of Preventive Medicine. The authors declare no conflict of interest.

## Author contributions

YK obtained funding, conceived, and supervised the project. DM, AK, SM, TC, JJP, and HM designed and performed experiments. FL analysed mass spectrometry data. TM designed CRISPR guides. DM and YK wrote the manuscript with input from all authors.

## Materials and Methods

### Cell culture

HEK293 and commercially available Flp-In T-REx HEK293 cells (Invitrogen; R78007) were cultured in DMEM (GIBCO) supplemented with 10% v/v Fetal Bovine Serum (FBS), 50μg/ml Penicillin Streptomycin, and 2mM L-Glutamine. Cell cultures were maintained in a 5% CO_2_ incubator in a humidified environment and routinely checked for mycoplasma. Cell lines were sourced from a dedicated facility at MRC-PPU core services.

### Cell fractionation

HEK293T cells (10 confluent plates) were collected in phosphate buffered saline/PBS (Gibco; 14190-094) supplemented with 1mM EDTA and 1mM EGTA. Cells were washed once in PBS, resuspended in ice cold cracking buffer (50mM Tris pH7.5, 1mM DTT, 0.1mM EDTA, 0.1mM EGTA), and incubated on ice for 15 minutes prior to lysis by mechanical stress (>20 sequential passes through 21-23-gauge needles). Lysate was cleared by centrifugation (17000xG for 5 minutes), passed through a 25mm/45μm polyethersulfone filter (Sigma Aldrich; WHA68962504), and de-salted into de-salting buffer (30mM MOPS pH7.0, 5% glycerol, 1mM DTT, 0.015% Brij 35) on a Sephadex G25 column using an Akta Pure Fast Protein Liquid Chromatography (FPLC) device. Lysate was next applied to a 1m Heparin HiTrap column (GE-Healthcare Life Sciences, now Cytiva) with elution on an increasing salt gradient into 1ml fractions (Greiner bio-one 96 well block; 780270). A salt gradient was introduced using a high salt buffer (30mM MOPS pH7.0, 1.2M NaCl, 5% glycerol, 1mM DTT, 0.015% Brij 35). Heparin column flow through was next passed through a Source Q HR 5/5 column into 30mM Tris-HCl pH8.2, 5% Glycerol, 1mM DTT, and 0.015% Brij 35 with elution on a salt gradient (High salt buffer supplemented with 1.0M NaCl). SourceQ fractions (1ml volume) were eluted into a 96 well block at 1ml intervals. Heparin and SourceQ binding fractions were immediately snap-frozen in liquid nitrogen and stored at −80°C until use. For mass-spectrometry fractions were washed/buffer exchanged in an Amicon ultra centrifugal concentrator (Millipore; UFC500396) with 2ml 30mM Tris-HCl pH 7.5 containing 1mM TCEP, followed by 2ml 50mM triethylammonium bicarbonate buffer (Sigma Aldrich T7408-100ml) containing 1mM TCEP. Samples were then concentrated to approximately 100μl volume and alkylated and with 40mM IAA (Sigma Aldrich; I1149-5G) for 3 hours in the dark at room temperature. Samples were next reduced by adding 2mM DTT and incubating at room temperature for a further 15 minutes (Formedium; DTT100). After overnight digestion with 10μg/ml mass-spectrometry grade Trypsin (Pierce; 90057), samples were submitted to the MRC-PPU mass-spectrometry facility for analysis.

### Western blotting

To prepare samples a 3:4 dilution was made in NuPAGE LDS sample buffer (ThermoFisher; NP007) supplemented with 10% v/v β-mercaptoethanol. Samples were heated to 95°C for 5 minutes prior to gel loading. Gel electrophoresis was performed using a XCell *SureLock* electrophoresis tank (ThermoFisher; EI0001) with 4-12% Bis-Tris NuPAGE pre-cast 12, 15 and 26 well gels (Invitrogen; NP0322, NP0323, and WG1403BX10). Protein was transferred to 45μm nitrocellulose membrane (Amersham; 10600002) at 90V for 90minutes in a Mini Trans-Blot Cell (Biorad;). Membranes were blocked for one hour in 5% Bovine Serum Albumin (Sigma Aldrich; A7906-100G)) dissolved in TBST. Primary antibodies were diluted 1:1000 in 5%BSA TBST and incubated overnight with shaking at 4°C. Membranes were washed three times for 10 minutes per wash in TBST. Membranes were next incubated with fluorescent secondary antibody (IR800) diluted 1:20,000 in TBS containing 5%BSA, 0.1% Tween for 30 minutes at RT with shaking. After washing in TBST for a further 30 minutes (3x 10-minute washes) membranes were visualised on a Lycor Odyssey CLx.

### Cell lysates

To generate cell lysates for RPL26 immunoblots (Fig-4), one near-confluent 15cm plate of HEK293 cells were gently collected in 0.5mM EDTA/0.5mM EGTA and placed on ice. Cells were washed once in ice cold PBS, and resuspended in lysis buffer (1%NP40, 50mM Tris-HCl pH7.5, 150mM NaCl) supplemented with protease inhibitor cocktail (1mM benzamidine, 1mM AEBSF, 1xprotease inhibitor cocktail (Roche; 48679800)). Lysates were cleared by centrifugation (20,000xG, 5 minutes) and mixed with LDS sample buffer as before. For enzymatic assays (Fig-1), cell lysates were generated in the absence of protease inhibitors as described in the cell fractionation procedure above. To induce ribosome stalling and UFMylation, cells were treated with 200nM Anisomycin dissolved in DMSO for 20 minutes prior to harvesting. Cytosolic and membrane fractions were isolated using digitonin and 1% NP40-based lysis buffers, respectively.

### Cloning

Pro-UFM1 (NP_057701.1) was cloned in frame with a N-terminal GFP tag and intervening short peptide linker (GSGEGRG) into the pcDNA5/FRT/TO bacterial expression vector (Invitrogen; V652020). A C-terminal histidine tag (Hisx6) with C3 protease site facilitated the generation of native protein after purification with NiNTA affinity beads. UFSP1 short, and long variant isoforms were cloned into the pGEX6P1 vector for bacterial expression by MRC reagents and services. UFSP2 (modified with a stabilising R136A mutation) was cloned into the petDuet (His6-TEV-UFSP2; DU59927) bacterial expression vector in frame with a N-terminal 6xHis-tag by Dundee University Reagents and Services division. A full list of cDNA constructs is included in Table-S1.

### Recombinant protein expression and purification

Human recombinant UCH-L1, UCH-L3, UCHL-5, and BAP enzymes were generated by MRC-PPU Reagents and Services. UFSP1 short (Q6NVU6) and long (A0A5F9ZGY7) isoforms were purified using standard Glutathione S-transferase (GST) affinity purification. Briefly, plasmids were transformed into *E.coli* BL21(DE3) competent cells and expression of the recombinant protein induced with 0.25M IPTG overnight (~16hours) at 18°C. Cells were sedimented by centrifugation, resuspended in lysis buffer (50mM Tris-HCl pH8.0, 300mM NaCl, 2mM DTT) supplemented with protease inhibitor cocktail (1mM benzamidine, 1mM AEBSF, 1xprotease inhibitor cocktail (Roche; 48679800)), and lysed by Ultrasonification. Lysate was cleared by ultracentrifugation at 30,000xg for 30 minutes and mixed with equilibrated Glutathione Sepharose 4B beads for approximately 1.5 hours at 4°C. Beads were washed twice in 250ml high salt wash buffer (50mM Tris-HCl pH8.0, 500mM NaCl, 2mM DTT) and once in 250ml low salt wash buffer (50mM Tris-HCl pH8.0, 150mM NaCl, 2mM DTT, 10%Glycerol). Protein was eluted by incubation with 0.1mg 3C protease (MRC-PPU Reagents and Services) in 10ml low salt wash buffer overnight at 4°C. Eluted protein was confirmed by SDS-PAGE gel electrophoresis and subsequently purified by gel filtration on a Superdex gel filtration column using FPLC. The prufied protein was aliquoted, snap-frozen in liquid nitrogen and stored at −80°C until further use.

### Expression and purification of recombinant UFM1-GFP fusion protein

pet15b-His-C3-UFM1-GSGEGR was transformed into BL21 competent E-coli (MRC-PPU Reagents and services division). Expression of the recombinant protein and cell lysis was performed as described above. Lysate was cleared by ultracentrifugation at 30,000g for 30 minutes, and then mixed with pre-equilibrated Ni^2+^ NTA beads in binding buffer (25 mM Tris 8, 300 mM NaCl, 10 mM Imidazole 2 mM DTT). Beads were then washed with 30 bed volumes of wash buffer (25 mM Tris 8, 300 mM NaCl, 20 mM Imidazole, 2 mM DTT). UFM1-GFP was eluted by incubation with 3C protease (MRC-PPU Reagents and Services) overnight at 4°C with occasional agitation. Gel filtration clean-up was performed with a Superdex 200 Hiload™ 16/600pg column on an Akta Pure FPLC device. Stocks were quantified as before, concentrated to 16mg/ml and immediately snap frozen in liquid nitrogen and stored at −80°C until use.

### GFP cleavage/DUB assay

Whole cell lysates (WCL) were next extracted from *UFSP2*^−/−^ HEK293 cells by mechanical lysis (syringe), thiol proteases ‘activated’ by addition of 10mM Dithiothreitol (DTT) (Kulathu *et al.*, 2013; Lee *et al.*, 2013) and incubated with the UFM1-GFP fusion protein. Cell fractions and/or recombinant enzyme were pre-activated on ice in Activation Buffer (50mM Tris-HCl pH7.5, 50mM NaCl) supplemented with 10mM freshly prepared DTT. Activated enzyme was next incubated with 5ug (3μM) recombinant UFM1-GFP fusion protein for 3 hours at 37°C. Cleavage of the GFP tag was analysed by Coomassie stain and/or immunoblot analysis. For assays involving chemical inhibitors, Iodoacetmide (Sigma-Aldrich; I1149-5G) or N-ethylmaleimide (Sigma-Aldrich; 04259-5G) were added to enzyme for 1 hour at room temperature in the dark prior to mixing with recombinant UFM1-GFP. All enzymatic reactions were completed in ultra-pure distilled water (Millipore QPOD; ZMQSP0D01).

### CRISPR-Cas9

CRISPR guide RNAs were designed with support from T. MacCartney at MRCPPU Reagents and Services. CRISPR sense and anti-sense guides were cloned into pX335 (Addgene plasmid 42335; Feng Zhang lab; Massachusetts Institute of Technology) and pBABED puro U6 (DU48788) plasmids respectively. The pX335 construct contains a chicken β–actin promoter-driven expression cassette for Cas9. In a separate strategy, single guide RNAs were cloned into the px459 vector (Addgene; 48139). Single guide-RNAs were targeted as follows; Exon-1 of UFSP1, Exon-5 of UFSP2. Full details of guide-RNAs, frameshift mutations and relevant sequencing data are included in the supplementary figures and key resources table. Procedures are described elsewhere (Ran, Hsu, Lin, *et al.*, 2013; Ran, Hsu, Wright, *et al.*, 2013); briefly, 1-2 million cells were seeded into a 10cm dish in antibiotic free Dulbecco’s Modified Eagle Medium (DMEM) and transfected with 1μg plasmid DNA using Lipofectamine 2000 (Invitrogen; 1168019) according to the manufacturer’s instructions. Cells were selected in 2μg/ml puromycin for 24 hours followed by a 24-hour recovery period in pre-conditioned media. Cells were plated at clonal dilution (0.7 cells/well) or submitted for single cell sorting, expanded, and screened for knockout by sequencing and/or immunoblot analysis.

### Sequencing

For UFSP1 and UFSP2 knockout clones, a ~1-1.5Kb fragment that included guide-RNA target sites was PCR amplified using Q5 High-Fidelity DNA Polymerase (NEB; M0491). Primers were designed using the NCBI Primer Blast tool and are documented in the key resources table. PCR products were purified by spin column (QIAGEN;28104) and cloned into a plasmid vector using the StrataClone blunt PCR cloning kit (Agilent; 240207). Colonies were selected and grown in 4ml 2xTY media supplemented with Ampicillin (10μg/ml). Plasmid DNA was extracted using the QIAprep Spin Miniprep kit (QIAGEN; 27104) and submitted for sequencing at the MRC PPU DNA sequencing and services division. Mutations were aligned to the Hg38 assembly (UCSC genome browser) using ClustalW (European Bioinformatics Institute; Muscle). Primers are detailed in Table-S2.

### Antibodies

primary antibodies; anti-UFM1 (Abcam; ab109305); anti-UCH-L1 (CusaBio; CsB-PA004381); anti-UFSP2 (Abcam; ab192597); anti-UFSP1 (Sigma-Aldrich; HPA027099); anti-GFP (Abcam; ab290); anti-RPL26 (Bethyl; A300-686A-M); anti-GAPDH (CST; 2118S); anti-HA (MRC-PPU reagents and services; 12CA5). Secondary antibodies; IRDye800 Donkey anti-Rabbit (Licor; 926-32213); IRDye800 Donkey anti-Mouse (Licor; 926-32212).

### Data visualisation and software

Western blots were processed in ImageStudio Lite (Licor) and arranged in Adobe illustrator. Original Graphics and cartoons were developed in Adobe Illustrator. Data filtering and analysis of public resources (MEROPS, GTEx) was completed in RStudio. Heatmaps were generated using the Complex Heatmap R-package (ComplexHeatmap) in R studio and clustered using default parameters (Euclidean method). Protein structures were visualised in ChimeraX.

### External resources

protein homology was analysed using Consurf (Tel Aviv University; https://consurf.tau.ac.il/) and visualised in Pymol (Educational license V2) by Schrodinger (https://pymol.org/2/). Mass-spectrometry results were aligned with the MEROPS database (European bioinformatics Institute; https://www.ebi.ac.uk/merops/) to discover novel peptidases. Gene expression data was downloaded from the Genotype-Gene expression project (GTEx) (Broad Institute; https://www.gtexportal.org/home/).

### Multiple sequence alignment

fasta files corresponding to the amino acid sequence of human UFSP1 (Q6NVU6; A0A5F9ZGY7) and UFSP2 (H0Y9B0; H0YA18; D6RA67; Q9NUQ7) protein coding transcripts were downloaded from Uniprot with reference to Ensembl annotation. Analagous sequences from other species were obtained with reference to the NCBI Homologene resource. Sequence alignment was performed using the ClustalW algorithm in Muscle (European Bioinformatics Institute). The ClustalW output was visualised in Jalview and edited in Adobe Illustrator.

### Data Independent Acquisition (DIA) proteomics

For each sample, a confluent 15cm plate of HEK293 cells was re-suspended in ice cold PBS (1mM EDTA/1mM EGTA), pelleted by centrifugation, and immediately lysed by addition of SDS-lysis buffer (5% SDS, 50mM TEAB pH8.5). lysates were boiled for 5 minutes at 95℃ followed by sonication using a Diagenode Biorupter at high energy for 10 cycles (30sec ON, 30sec OFF). Lysates were cleared by centrifugation at 20,000xG for 20 minutes and quantified by BCA assay (Pierce; 23225). 200μg protein was prepared as follows; TCEP stock solution (100mM TCEP, 300mM TEABC) was added to final concentration of 10mM TCEP (1:10) and samples incubated at 60℃ for 30 minutes. Samples were rested at room temperature and freshly prepared iodoacetamide (IAA) added to 40mM final concentration. After 30 minutes at room temperature shielded from light and with gentle agitation, samples were acidified by addition of mass-spectrometry grade 12% phosphoric acid to a final concentration of 1.2% (1:10). Sample ‘clean-up’ was completed using S-trap micro-columns with an overnight on-column digestion using 13μg trypsin per 200μg of protein input. Eluted peptides were lyophilised by speed-vacuum and submitted to the MRC-PPU core mass-spectrometry facility. For differential expression analysis, data was processed using the POMA R-package and LIMMA. To identify modified lysine residue DIA data was searched for the KVG sequence (UFM1 C-terminal VG remnant) using Spectronaut 15. Selected MSMS spectra of VG modified peptides were annotated using IPSA (http://www.interactivepeptidespectralannotator.com).

---

**Figure-S1.**
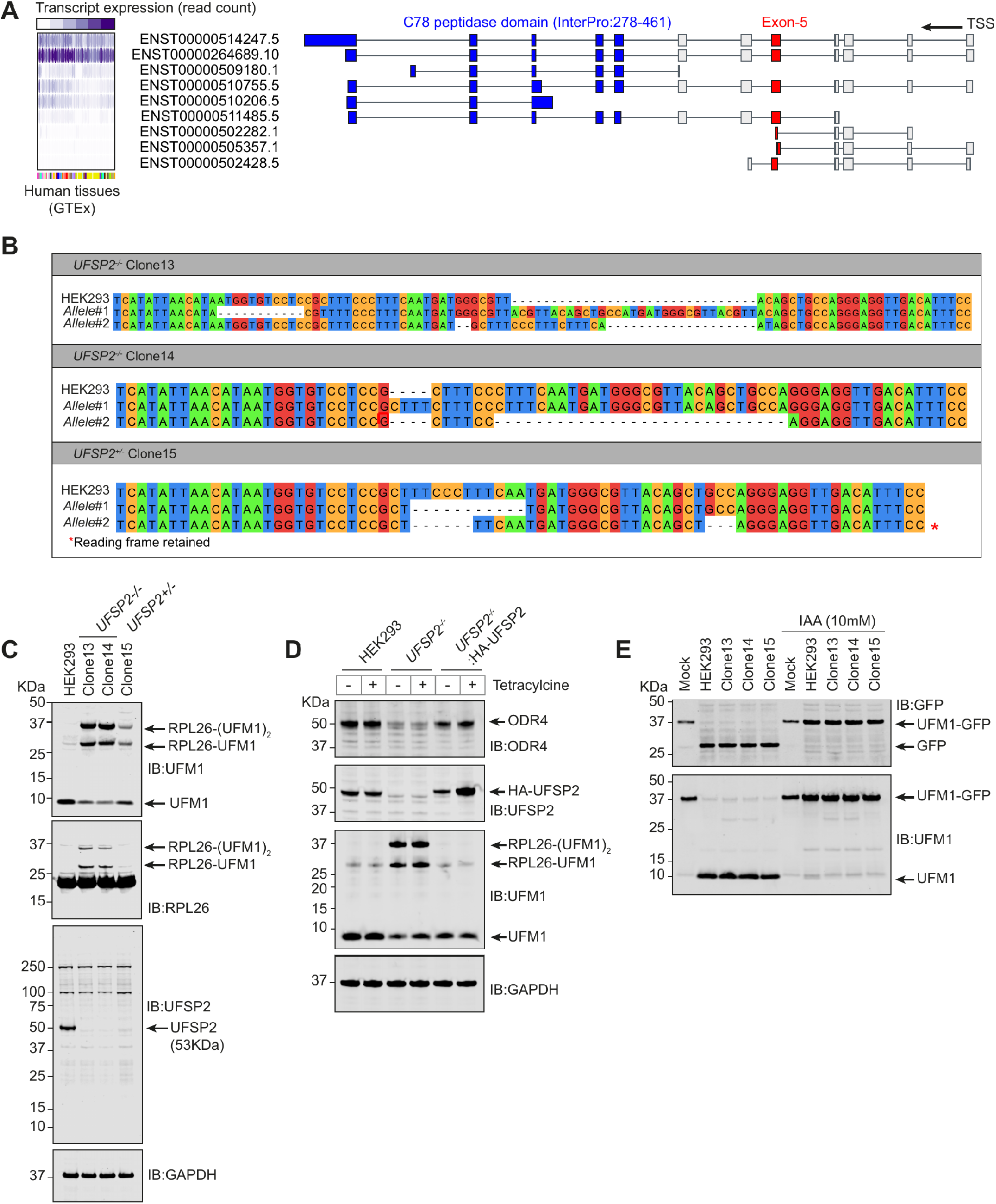
Targeted knockout of UFSP2. **(A)** Schematic of protein coding transcripts of UFSP2 as annotated by the GTEx resource (Broad Institute). Figure was downloaded from the GTEx transcript browser and edited in Adobe Illustrator. Regions coding for the C78 protease domain and CRISPR target region are shown in blue and red respectively. Gene expression data (read count) is condensed and shown left – consult https://gtexportal.org/home/ for full data. **(B)** Sequencing analyses for CRISPR clones designated 13, 14, and 15. Exon5 was PCR amplified and cloned using the StrataClone blunt ended cloning kit (Agilent). Sequences are representative of at least 8 bacterial colonies selected for miniprep and sequencing. Mutations were determined by multiple sequence alignment (ClustalW) to the Hg38 reference genome. Note that Clone 15 retains an active reading frame in one allele. Excerpt from the UCSC genome browser is shown top for reference **(C)** Immunoblot analysis of cell lysate from all clones. **(D)** Rescue of *UFSP2*^−/−^ cell line phenotype by stable transfection of UFSP2 expressing cDNA. The HEK293 Flp-in TREx cell line was transiently transfected with pcDNA5 expressing HA-tagged UFSP2 and subject to antibiotic selection. **(E)** Cleavage assay using cell lysates derived from all clones. Where indicated, lysates have been pre-treated with Iodoacetamide (IAA) for one hour at room temperature in the dark. Experiments in C-D are representative of more than three experiments.

**Figure-S2.**
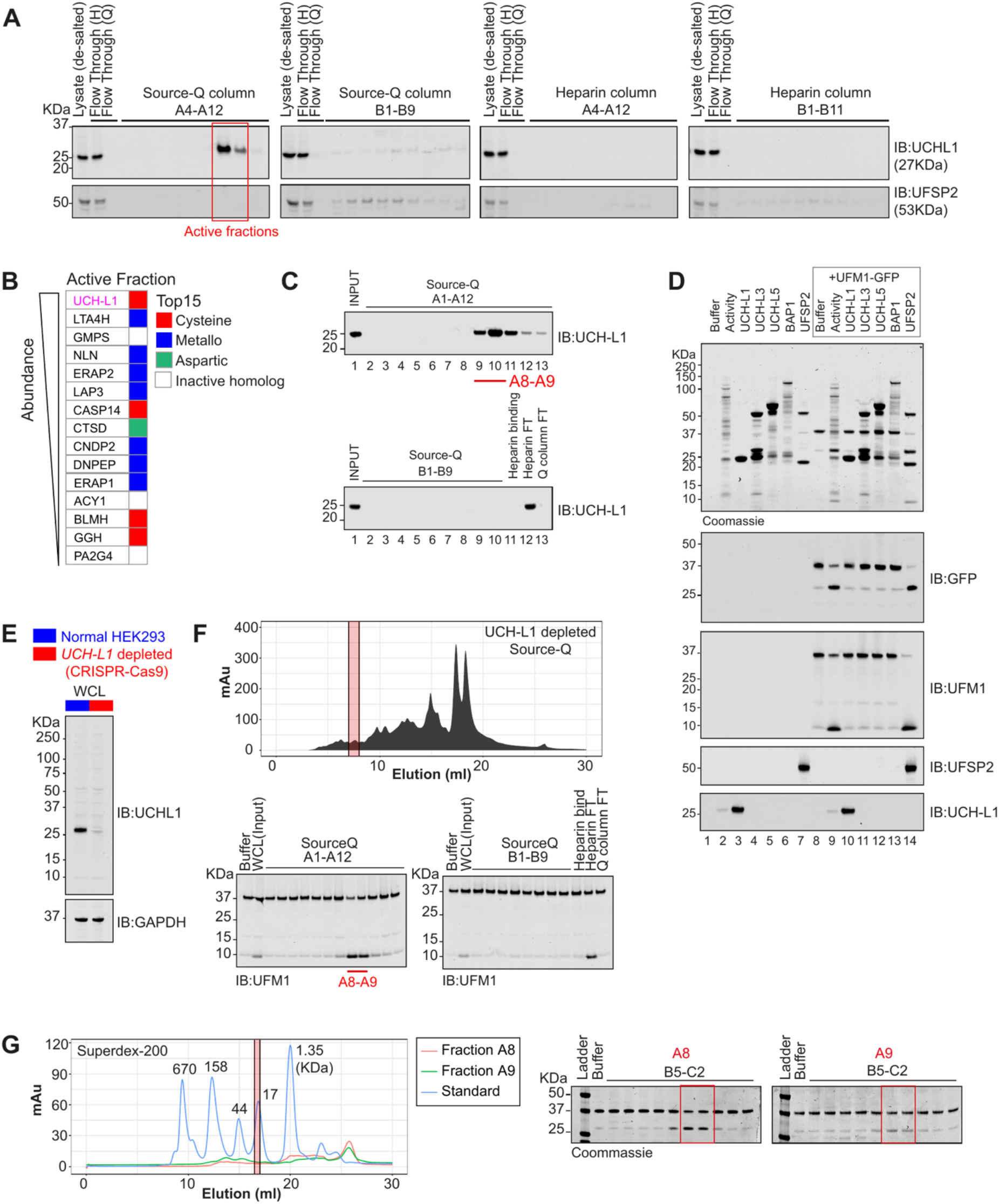
UCH-L1 depletion and bioinformatic characterisation of UFSP isoforms. UCH-L1 was identified as a top hit in our initial screens. Speculatively, we considered that UCH-L1 may obstruct detection of the novel peptidase either by sequestration of column volume or through the saturation of mass-spectrometry detection. Therefore, CRISPR-Cas9 was employed deplete UCH-L1 expression in HEK293 cells prior to the fractionation procedure. Data presented in Fig-2D-E is from experiments using UCH-L1 depleted cells. **(A)** Representative blotting of fractions eluted from Heparin and Source-Q columns showing UFSP2 distribution in relation to UCH-L1. Red boxes highlight the active fractions. **(B)** LC/MS analysis of active fractions derived from HEK293 cells showing UCH-L1 as the most frequently identified protease. Only the top15 proteases are shown. **(C)** Immunoblot analysis showing distribution of UCH-L1 in fractions. Experiment is same as shown Fig-2B. **(D)** In vitro cleavage assay; the UFM1-GFP probe incubated with UCH-L1 and related DUBs. **(E)** Immunoblot analysis of UCH-L1 depleted cell lines generated through transfection of *UCHL1* targeting CRISPR-Cas9 constructs. Small amounts of UCH-L1 are detected by LC/MS shown Fig-2D. **(F)** (top) Akta chromatogram showing fractionation of UCH-L1 depleted HEK293 cells (HiLoad^®^ 16/600) (below) *In vitro* cleavage assay incubating fractions with UFM1-GFP probe. **(G)** fraction A8 and A9 in were subject to further clean-up on a Superdex^®^ 200 pg column. Activity in fractions was confirmed by cleavage assay (right) prior to analysis by LC/MS. Resulting data is shown Fig-2D.

**Figure-S3.**
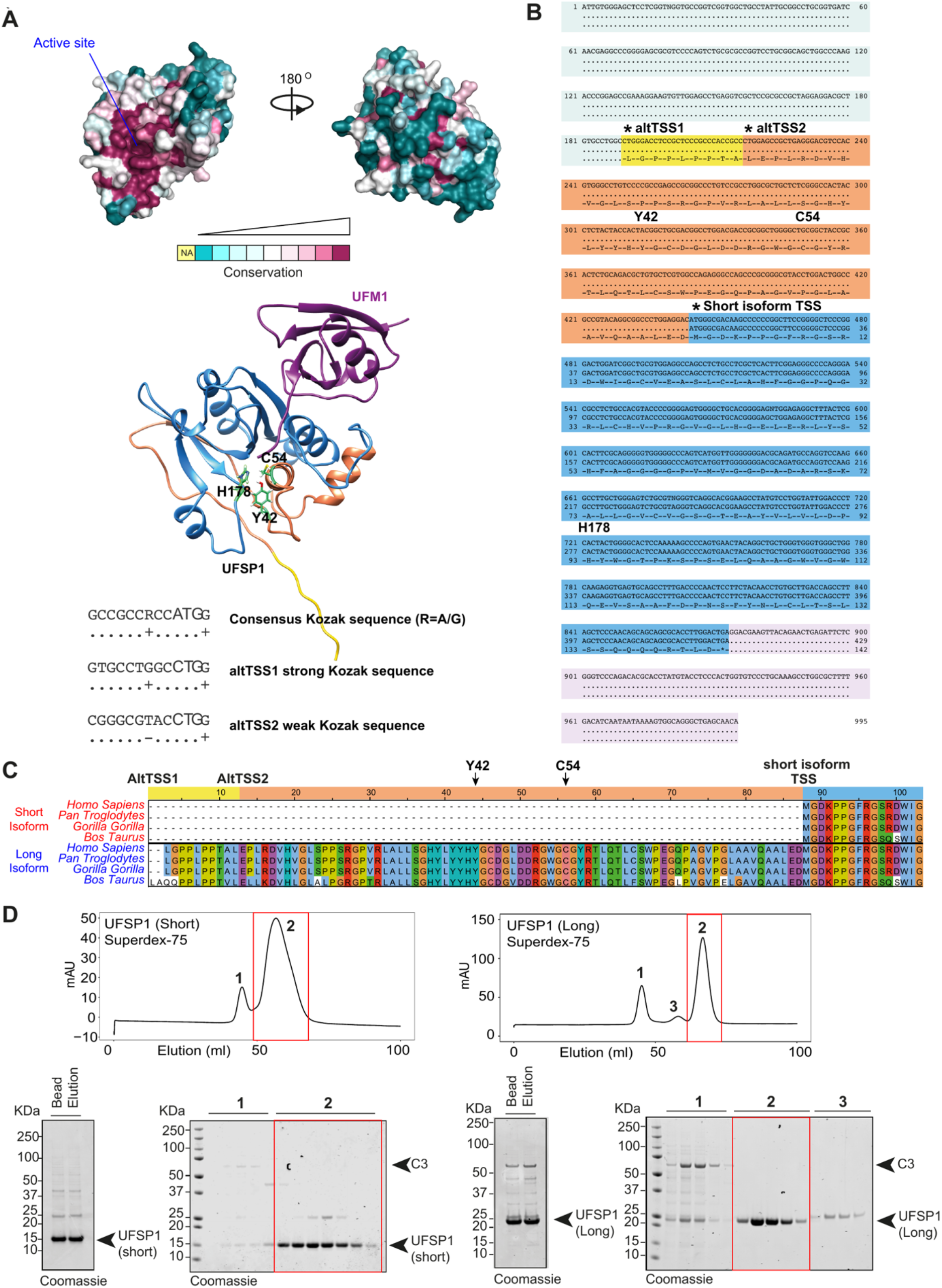
UFSP1 expression and purification. **(A)** (top) Consurf coloration of reported crystal structure of murine UFSP1 (pdb 2Z84). (below) Alphafold prediction of UFSP1 in complex with UFM1. Coloring corresponds to sequence analysis shown in (B). **(B)** Bioinformatic analysis of UFSP1 transcripts. Highlighted are key features including predicted transcription start sites and catalytic residues. **(C)** Cross-species multiple sequence alignment highlighting long isoform unique residues **(D)** Gel-filtration clean-up of human recombinant short (Q6NVU6) and long (A0A5F9ZGY7) UFSP1 isoforms. Raw elution from beads, after immunoprecipitation and 3C cleavage, is shown left for reference. Duplicate lanes are technical replicates.

**Figure-S4.**
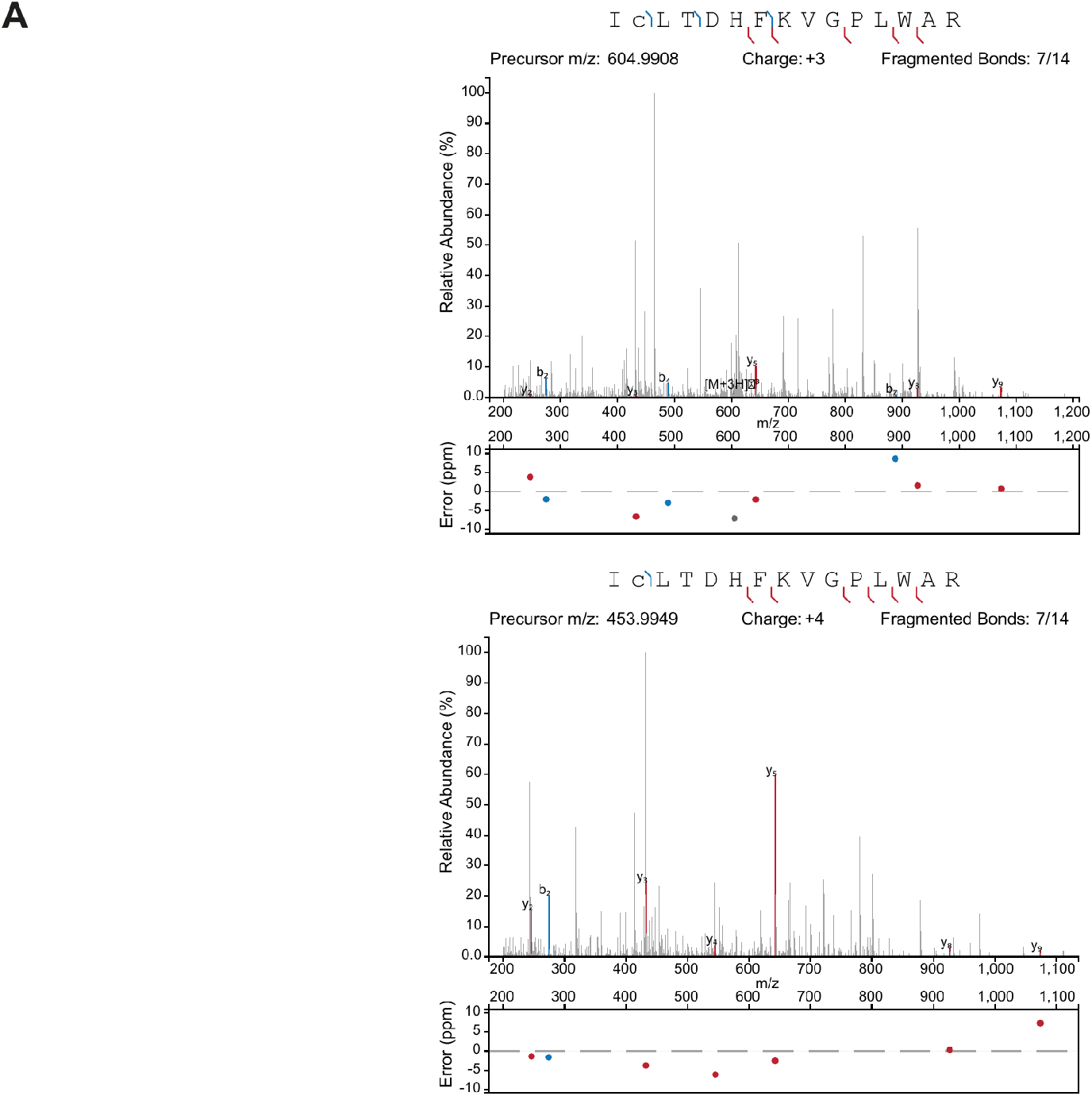
Modification of UFC1 Lysine 122 with UFM1. **(A)** MSMS spectra showing K122 joined to the expected VG motif resulting from UFMylation. Spectra were generated from the analysis of DIA proteomics using Spectronaut 15. Selected MSMS spectra of VG modified peptides were annotated using IPSA (http://www.interactivepeptidespectralannotator.com)(Brademan *et al.*, 2019).

**Figure-S5.**
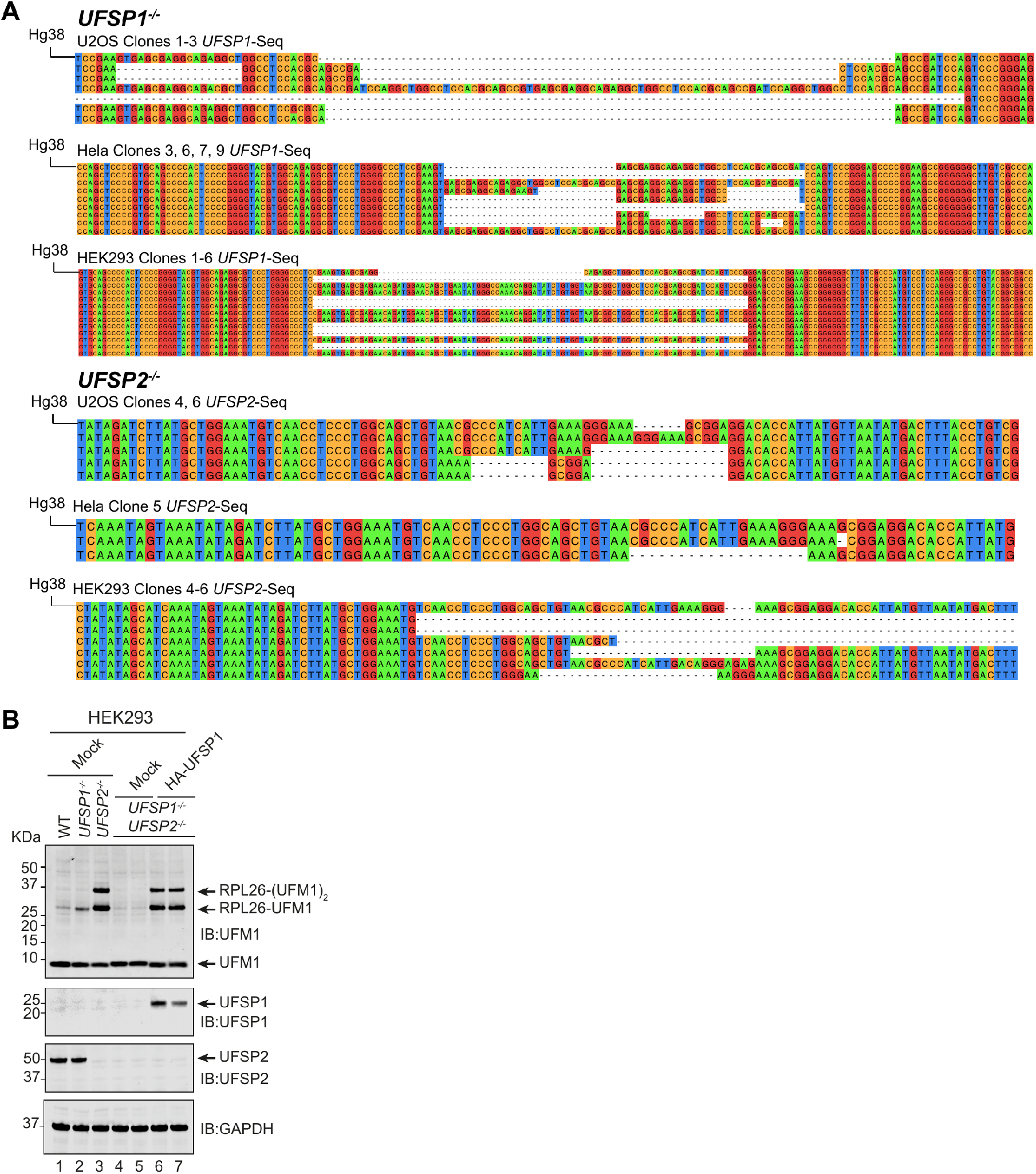
UFSP1 function in ribosome UFMylation. **(A)** Representative sequencing data for clones shown in Fig-5A. Each clone was sequenced a total of 8 times and aligned with the reference genome (Hg38) using Muscle (European Bioinformatics Institute). Shown are two sequencing traces representative of each allele, at the approximate location of Cas9-induced mutation. Mutations were determined by multiple sequence alignment (ClustalW) of sequencing traces to the Hg38 reference genome. **(B)** Rescue of loss-of-function phenotype by UFSP1 over-expression. *UFSP1*^−/−^/*UFSP2*^−/−^ HEK293 cells were transiently transfected as indicated. The anti-UFM1 western blot is reproduced in Fig-5D.

**Figure-S6.**
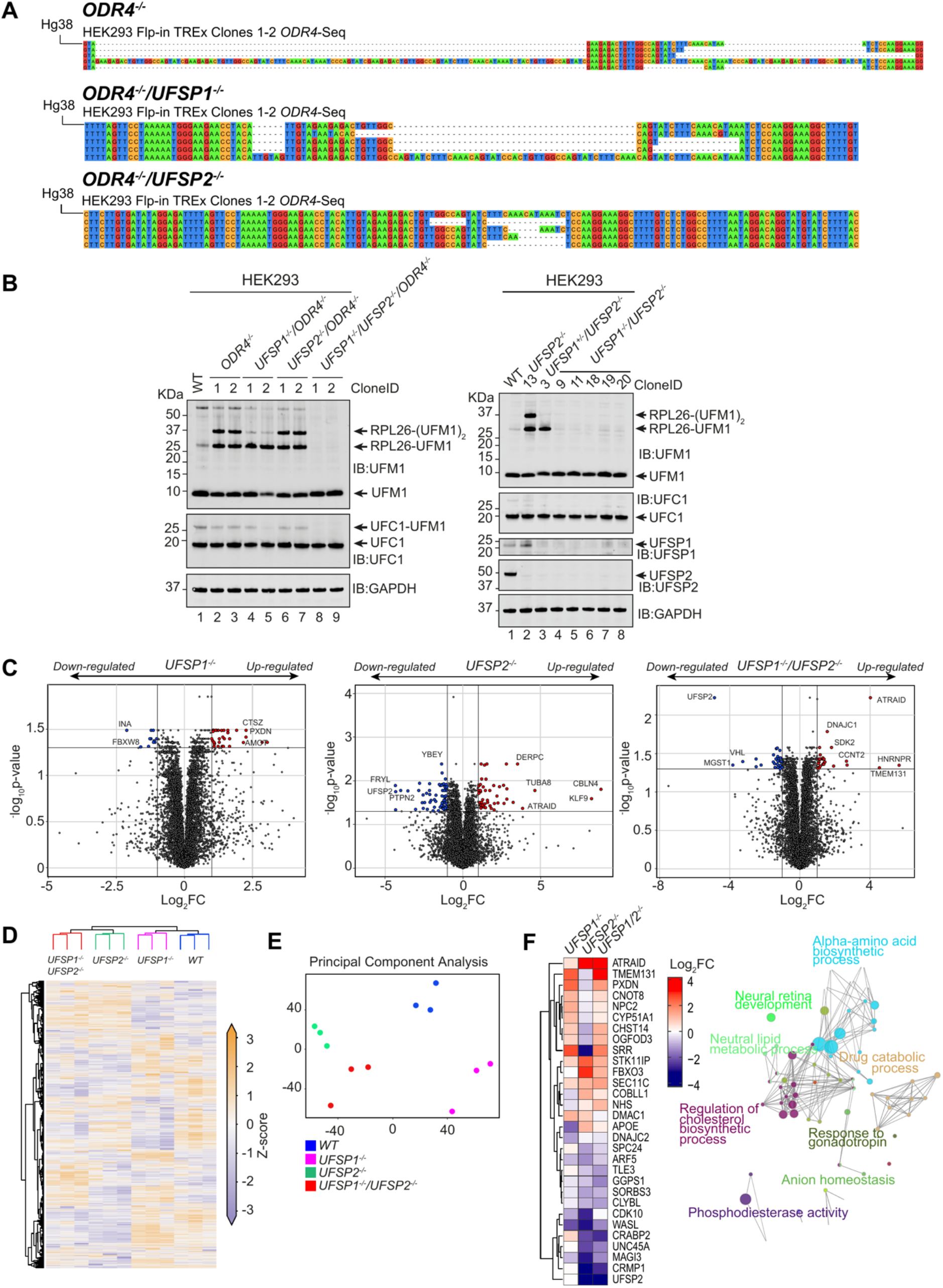
ODR4 function in UFMylation. **(A)** Representative sequencing analysis of the *ODR4* locus. Shown are two alleles identified from 8 sequencing reactions aligned to the Hg38 reference genome. Clone-ID (MRC-PPU internal) for analysis in Fig-6A is as follows; *UFSP1*^−/−^ (Clone-1); *UFSP2*^−/−^ (Clone-13; as in Fig-S1); *UFSP1*^−/−^ /*UFSP2*^−/−^ (Clone-11); *ODR4*^−/−^ (Clone-1); *ODR4*^−/−^/*UFSP1*^−/−^ (Clone-1); *ODR4*^−/−^/*UFSP2*^−/−^ (Clone-1). Full sequencing data is available on request from the study lead author. **(B)** Immunoblot analysis of the indicated clones. Clone-3 (D) is heterozygous deficient for *UFSP1* and shows a selective defect in the second UFM1 modification on RPL26 (K134). **(C)** Data Independent Acuisition (DIA) quantitative proteomics of indicated knockout cell lines. Volcano plots showing differential expression analysis (LIMMA). **(D)** (left) heatmap analysis of combined data (padj<0.05) (right) principal component analysis shows hierarchical clustering (Euclidean) of technical replicates. Technical replicates are different plates of the same clone processed in parallel. **(E)** Principal component analysis of proteomics data. Each dot of the same color is a technical replicate. **(F)** (left) Representative examples of heatmap shown in Fig-6C, selected at random using R-studio/dplyr package (sample_n=30) and hierarchically clustered by the Euclidean method; (right) network summarising GO-term enrichment resulting from genes passing statistical thresholds (shown Fig-6C). Generated in Cytoscape using the ClueGO app (hypergeometric test).

